# Antisense oligonucleotide-mediated correction of *CFTR* splicing improves chloride secretion in cystic fibrosis patient-derived bronchial epithelial cells

**DOI:** 10.1101/2020.05.12.089417

**Authors:** Wren E. Michaels, Robert J. Bridges, Michelle L. Hastings

## Abstract

Cystic fibrosis (CF) is an autosomal recessive disorder caused by mutations in the CF transmembrane conductance regulator (*CFTR*) gene, encoding an anion channel that conducts chloride and bicarbonate across epithelial membranes. Mutations that disrupt pre-mRNA splicing occur in more than 15% of CF cases. One common *CFTR* splicing mutation is *CFTR* c.3718-2477C>T (3849+10kbC>T), which creates a new 5’ splice site, resulting in splicing to a cryptic exon with a premature termination codon. Splice-switching antisense oligonucleotides (ASOs) have emerged as an effective therapeutic strategy to block aberrant splicing. We test an ASO targeting the *CFTR* c.3718-2477C>T mutation and show that it effectively blocks aberrant splicing in primary bronchial epithelial (hBE) cells from CF patients with the mutation. ASO treatment results in long-term improvement in CFTR activity in hBE cells, as demonstrated by a recovery of chloride secretion and apical membrane conductance. We also show that the ASO is more effective at recovering chloride secretion in our assay than ivacaftor, the potentiator treatment currently available to these patients. Our findings demonstrate the utility of ASOs in correcting CFTR expression and channel activity in a manner expected to be therapeutic in patients.

## INTRODUCTION

Cystic fibrosis (CF) is a fatal, autosomal recessive disorder caused by mutations in the cystic fibrosis transmembrane conductance regulator (*CFTR*) gene (1). *CFTR* encodes a cAMP-regulated membrane channel that conducts chloride (Cl^-^) and other negatively charged ions, such as bicarbonate (HCO_3_-), across epithelial cell membranes (2). Mutations in *CFTR* alter epithelial anion secretion leading to disruption of ionic and water homeostasis in epithelial cells of a number of organs including lungs, pancreas, and intestine (3). Pharmacotherapy for CF has focused on the development of two main categories of drugs: correctors that improve the expression of CFTR, and potentiators that stimulate channel function (4, 5). These therapeutics have proven effective in treating a large proportion of CF patients with mutations that disrupt protein expression and channel function, such as the common F508del-*CFTR* (ΔF508) and G551D mutations (6–10). However, these types of therapeutics are less effective in treating patients with mutations that affect pre-mRNA splicing, as these types of mutations often result in the disruption of the *CFTR* reading frame.

*CFTR* splicing mutations account for ∼15% of known CF-associated mutations (11–13). *CFTR* c.3718-2477C>T (also known as *CFTR* 3849+10kb C>T and c.3717+12191C>T) is the 10^th^ most common *CFTR* mutation worldwide (CFTR2: www.cftr2.org). The C>T change generates a de novo donor 5’ splice site downstream of exon 22 in intron 22 that activates a new 84-nucleotide pseudo-exon (referred to here as ψ-Ex) containing a premature termination codon (Figure 1A)(12, 14). Splicing to the cryptic (ψ-Ex) exon is variable, and ranges from 0-92% of total *CFTR* mRNA from the mutated allele (14, 15). Disease severity in patients with the mutation is also variable and previous studies on the *CFTR* c.3718-2477C>T mutation have shown a correlation between the amount of aberrant splicing and disease severity (14–18). This correlation suggests that even a partial block in splicing of the cryptic splice site and an increase in normal splicing could be beneficial to CF patients.

**Figure 1.**
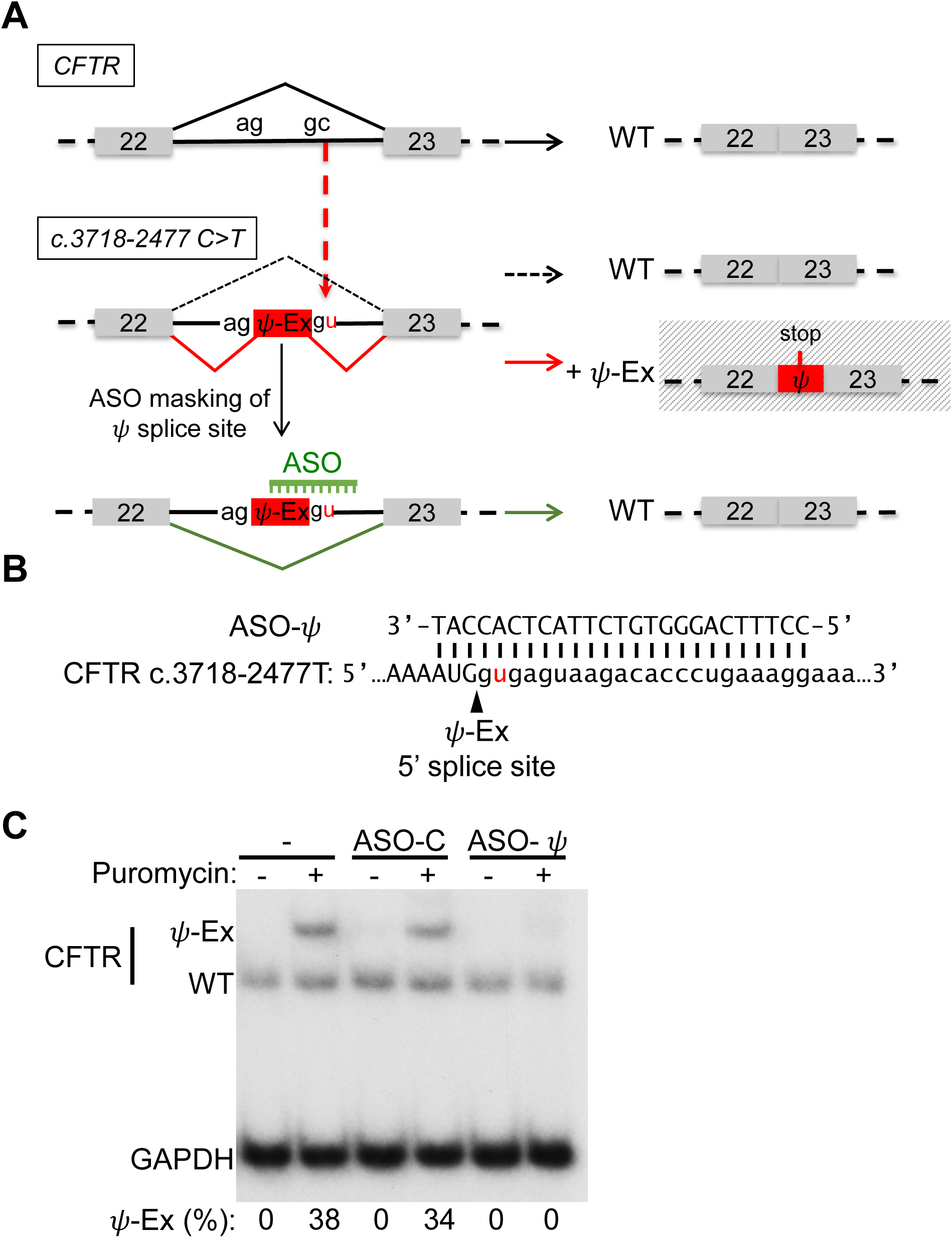
*CFTR* c.3718-2477C>T aberrant splicing and correction with a splice switching antisense oligonucleotide. (**A**) Schematic of the aberrant splicing caused by the *CFTR* c.3718-2477C>T mutation in intron 22. The C>T mutation creates a de novo 5’ splice site resulting in the insertion into the mRNA of an 84 nucleotide pseudoexon (ψ-Ex) that has an in-frame stop codon. An ASO designed to mask the *CFTR* c.3718-2477C>T de novo splice site redirects splicing to the canonical splice site, restoring wild-type (WT) mRNA. (**B**) Sequence alignment of ASO-ψ to the ψ-Ex 5’ splice site found in *CFTR* c.3718-2477C>T. (**C**) RT-PCR analysis of *CFTR* splicing in a lymphoblast cell line from a CF patient homozygous for the *CFTR* c.3718-2477C>T mutation following transfection with vehicle only (-), a control, non-targeted ASO (ASO-C) or ASO-ψ targeting the ψ-Ex 5’ splice site. Cells were untreated (-) or treated (+) with puromycin before RNA isolation. *GAPDH* was analyzed as a control for total cellular RNA abundance. Splicing was quantified as the percent of RNA transcripts with ψ-Ex [(ψ-Ex/(ψ-Ex+WT))x100] and is shown below each lane.

Splice-switching antisense oligonucleotides (ASOs) have proven to be effective therapeutic modulators of pre-mRNA splicing in a target-specific manner (19–21). ASOs are short, chemically modified oligonucleotides that bind to a target sequence through complementary, anti-parallel base-pairing. ASOs designed to alter splicing act by base-pairing to the target RNA and sterically interfering with splice site recognition by splicing factors (20, 22).

ASOs have been explored previously as a tool to correct defective splicing in CF (11, 23–26). For the *CFTR* c.3718-2477 splicing mutation, an ASO that blocks the cryptic splice site was tested in a CFTR expressing mini-gene system that partially recapitulated the cryptic splicing associated with the mutation and the results showed efficient ASO-mediated recovery of correct splicing and consequent elevation in CFTR protein (25). This study was an important first demonstration of an ASO-based correction of *CFTR* c.3718-2477T RNA splicing.

In the current report, we provide a critical next step in the testing of an ASO-based therapeutic candidate for this mutation by demonstrating that treatment of patient-derived cells with an ASO that masks the cryptic splice site restores splicing at the correct splice site and improves CFTR channel function. This improvement in CFTR channel function is comparable to that observed with C18 (VX-809 analog) and VX-770 treatment, current FDA approved drugs (VX-809 and VX-770) for the treatment of F508del-*CFTR* patients, and superior to these drugs in the treatment of patients with the *CFTR* c.3718-2477C>T mutation (27, 28). Our results support the use of ASOs for the treatment of patients with the *CFTR* c.3718-2477C>T mutation and provides a basis for advancing ASOs for the treatment of patients with other *CFTR* splicing mutations.

## MATERIAL AND METHODS

### Nomenclature

For *CFTR* mutations, the HGVS cDNA name is used. The legacy name (CFTR2: www.cftr2.org) is provided in parentheses after the first citation within the text (29).

### Cells and culture conditions

The human patient lymphoblast cell line (GM11860; Coriell Institute) was grown in RPMI media supplemented with 15% Fetal Bovine Serum (FBS). Primary bronchial human epithelial (hBE) cells were obtained under Institutional Review Board approved protocols at the University of North Carolina and University of Pittsburgh. In brief, CF hBE cells were isolated from patient lung explants and expanded in culture in hBE growth media using established protocols (30). Cultures were from two donors compound heterozygous for the *CFTR* c.3718-2477C>T splice site mutation and F508del-*CFTR* (Patient codes KKD004M, donor 1, and KKCFFT021K, donor 2), one donor homozygous for the splice mutation (Patient code KK018E, donor 3), two non-CF donors with WT-*CFTR* (Patient codes HBE0125, donor 5 and HBE069, donor 6), and one donor homozygous for F508del-*CFTR* (Patient code BRCF001, donor 4). The hBE cells were differentiated by plating on Costar 24-well high-throughput screening (HTS) filter plates (0.4 μM pore size, Polyester, Corning, catalog #CLS3397) and growing at an air-liquid interface in hBE differentiation media containing Ultroser-G (2%) in a 37°C incubator with 90% humidity and 5% CO_2_ for 5-7 weeks. Media was replaced on the basolateral side 3 times a week.

### Antisense oligonucleotides

Splice-switching antisense oligonucleotides are 25-mer phosphorodiamidate morpholino oligomers (PMO) Gene-Tools, LLC (Supplementary Table S1). A non-targeting PMO was used as a negative control, ASO-C, (Gene Tools, standard control oligo). ASOs were formulated in sterile water.

### Cell transfection

ASOs were transfected at a final concentration of 15 μM in lymphoblast cell lines and 40 μM in hBE cells using Endo-Porter according to manufacturer instructions (Gene Tools, LLC)(31). Lymphoblast RNA was collected 48 hours post-transfection. Primary hBE cells were transfected after differentiation on filter plates. For this, the mucus layer was removed four days before transfection by incubating the apical surface in 70 μL of PBS supplemented with 3 mM DTT for 30 minutes followed by aspiration. The apical surface was similarly washed with PBS the day before transfection. Cells were transfected with ASOs apically in 100 μL of a hypo-osmotic solution [1:1, Dulbecco’s Phosphate-Buffered Saline (DPBS):dH_2_O] for one hour at which time the solution was removed and the cells were re-transfected with the same ASO in 100 μL of DPBS at a final concentration of 40 μM (32). The solution was fully absorbed at the apical surface after four days, at which time the chloride channel function was assessed as described below. For the time-course study, PBS alone (100 μL) was added apically every 4 days post-transfection. Twelve hours after the transfection of ASO, lymphoblast cells were treated for nine hours with puromycin at a final concentration of 200 μg/ml, after which time RNA was collected (33). Differentiated hBE cell cultures were treated apically with 200 μg/ml of puromycin for 6 hours immediately following transepithelial current clamp (TECC) analysis in HEPES-buffered Coon’s F12 media and immediately before RNA isolation.

### RNA isolation and RT-PCR

RNA was extracted from cells using TRIzol according to manufacturer instructions (Thermo Fisher Scientific). RNA was extracted from differentiated cells grown on filters by cutting out the filter and placing directly in TRIzol. Reverse transcription was performed on total RNA using the GoScript Reverse Transcription System with an oligo-dT primer (Promega). Splicing was analyzed by radiolabeled PCR of resulting cDNA using GoTaq Green (Promega) supplemented with α-^32^P-deoxycytidine triphosphate (dCTP). Primers for amplification are reported in Supplementary Table S1 and include primer sets flanking human *CFTR* exons 22 and 24 (hCFTRex22F, hCFTRex24R), and primers for human GAPDH, human β-actin (hβ-actinFor, hβ-actinRev) and SRSF2 1.7kb (hSC351.7kbF, hSC351.7kbR)(34). Reaction products were run on a 6% non-denaturing polyacrylamide gel and quantified using a Typhoon 7000 phosphorimager (GE Healthcare).

### Real-time qPCR

Real-time qPCR was performed with PrimeTime Gene Expression Master Mix and PrimeTime qPCR probe assay kits targeting the 84 base-pair insert (hCFTR-2477), non-F508del-*CFTR* (hCFTR-F508), and F508del-*CFTR* (hCFTR-DF508) transcripts normalized to human β-actin (IDT, HsPT.39a.2214847)(Supplementary Table S1). All reactions were analyzed in triplicate on 96-well plates and averaged together to comprise one replicate. Real-time PCR was performed on an Applied Biosystems (ABI) ViiA 7 Real-Time PCR System with the thermal-cycling protocol: stage 1-50°C for 2 min, 95°C for 3 min; stage 2 – 40 cycles of 95°C for 15 s, 60°C for 1min to ensure an amplification plateau was reached. A TaqMan assay kit for human β-actin was utilized (Hs.PT.39a.22214847, IDT). Results were analyzed by the ΔΔCT method (35).

### Protein isolation and automated western analysis

Cell lysates for immunoblot analysis were prepared from patient hBE cells (donor 3: patient code KK018E) after transfection and functional analysis using NP-40 lysis buffer (1% Igepal, 150mM NaCl, 50mM Tris-HCl pH7.6) supplemented with 1× protease inhibitor cocktail (Sigma-Aldrich). Protein concentration was measured using a Coomassie (Bradford) protein assay. Cell lysates were prepared and diluted to 3 mg/ml using the sample preparation kit (Protein Simple) for an automated capillary western blot system, WES System (Protein Simple) (36, 37). Cell lysates were mixed with 0.1× sample buffer and 5× fluorescent master mix for a final concentration of 1.5 mg/ml. Samples were incubated at room temperature for 20 minutes and then combined with biotinylated protein size markers, primary anti-CFTR antibodies 570, 450 (Riordan lab UNC, Cystic Fibrosis Foundation, diluted 1:50;1:200 respectively with milk-free antibody diluent), anti-SNRPB2 (4G3, Krainer lab, Cold Spring Harbor Laboratory, diluted 1:2000 with milk-free antibody diluent), horseradish peroxidase (HRP)-conjugated secondary antibodies, chemiluminescence substrate and wash buffer and dispensed into respective wells of the assay plate and placed in WES apparatus. Samples were run in triplicate. A single outlier was identified using ROUT analysis (Q=1%) and removed. Signal intensity (area) of the protein was normalized to the peak area of the loading control SNRPB2, small nuclear ribonucleoprotein polypeptide B2. Quantitative analysis of the CFTR B and C-bands was performed using Compass software (Protein Simple).

### Automated equivalent current assay

Differentiated epithelial cells were treated three days post-transfection basolaterally with C18 (6 μM) (VRT-534, VX-809 analog) or vehicle (0.01% DMSO) at 37°C (38). Twenty-four hours later the cells were switched from differentiation media to HEPES-buffered (pH 7.4) F12 Coon’s modification media (Sigma, F6636) apically and basolaterally and allowed to equilibrate for one hour at 37°C without CO_2_. To obtain the calculated equivalent current (I_eq_) measurements, the transepithelial voltage (Vt) and resistance (Rt) were recorded at 37°C with a 24-channel TECC robotic system (EP Design, Belgium) as previously described (39). Briefly, baseline current measurements were taken for ∼40 minutes. Benzamil (6 μM final concentration) was then added to the apical side and measurements were taken for approximately 20 minutes. Forskolin (10 μM) was added to the apical and basolateral sides and Vt and Rt measurements were recorded for 20 minutes. The cells were then treated apically and basolaterally with potentiator, VX-770 (1 μM), and measurements were taken for an additional 20 minutes. The cells were then treated with the inhibitor bumetanide (20 μM) on the basolateral side for 30 minutes. Measurements were taken at two-minute intervals. I_eq_ was calculated using Ohm’s law (I_eq_ = Vt/Rt) and plotted as current traces. Area under the curve (AUC) measurements of forskolin and forskolin + VX-770 were calculated using a one-third trapezoidal rule for the entirety of each test period. The average of two identical wells was calculated for each plate. Each final replicate was an average of two plates (four wells total, n=4) treated simultaneously to obtain the final mean ±SEM graphed for each donor. A total of nine independent experimental groups (N=9; 36 wells total) were used for donor 1 (KKD004M), three (N=3; 12 wells) for donor 2 (KKCFFT012K) and four (N=4; 16 wells) for donor 3 (KK018E).

### Transepithelial impedance analysis

Impedance analysis was performed on hBE cells cultured on filters and treated as in the automated equivalent current studies, above, and as described previously (40, 41). Impedance was measured at 37°C at 126 frequencies. The spectra are presented as Nyquist plots and fit to equivalent circuit models of epithelial membranes to obtain separate apical and basolateral resistances. Membrane conductance was calculated as a reciprocal of resistance. Vt, Rt values and impedance spectra for 24 wells were obtained simultaneously. Baseline measurements were taken for 24 minutes. Benzamil (6 μM final concentration) was then added to the apical side. Changes in apical conductance were measured by the change in resistance upon stimulation with forskolin (10 μM) to the apical and basolateral sides following benzamil treatment. Subsequently, the cells were treated apically and basolaterally with potentiator, VX-770 (1 μM). After treatment with the VX-770, the cells were treated with two doses of CFTR inhibitor GlyH-101 (10 μM and then 60 μM) on the apical and basolateral sides. Measurements were taken for 10 minutes after each addition. Membrane conductance was calculated by fitting the impedance spectra to the equivalent electric circuit models of epithelial membranes as described previously using a program developed by the Bridges’ lab (40). Spectra and current measurements were taken five times at 2.5-minute intervals for each experimental period and the four calculated conductance measurements were averaged correlating to the four I_eq_ plateau measurements, excluding the initial peak immediately following the addition of forskolin. The average of two identical wells was calculated for each plate (n=2) to obtain one replicate. A total of two independent experimental plates were used for the final plotted calculated mean ±SEM graphed (N=2).

### Statistics

Statistical analyses were performed using GraphPad PRISM 8.2.1 or Microsoft Excel. A two-tailed one-sample t-test was used to assess significant changes of one test-group normalized to a control. One-way ANOVA analysis with a post-hoc test (Tukey’s) was used to assess significant differences when comparing more than two groups. Two-way ANOVA analysis was used when comparing two independent variables with a post-hoc test (Tukey’s) to assess significant differences between and within groups. When assessing significance differences within groups of paired data Sidak’s post-hoc test was used. The specific statistical test used in each experiment can be found in the figure legends.

## RESULTS

### ASO-ψ blocks ψ-Ex splicing in a *CFTR* c.3718-2477C>T lymphoblastoid patient cell line

As an initial analysis of an ASO designed to block splicing at the de novo donor 5’ splice site created by the *CFTR* c.3718-2477C>T mutation, we tested ASO-ψ (Figure 1B). ASO-ψ base-pairs to the donor splice site created by the mutation, a similar region that a previously reported ASO targeted (25), though we altered the length, chemistry, and transfection procedure in order to achieve effective delivery to patient-derived cells *in vitro*. ASO-ψ was transfected into a lymphoblastoid cell line derived from a patient homozygous for *CFTR* c.3718-2477C>T and splicing was analyzed by RT-PCR (Figure 1A, B). Initial analysis of RNA isolated from untreated cells revealed that only correctly spliced *CFTR* mRNA was detected, with no apparent splicing to the 84 base-pair pseudo-exon, ψ-Ex, associated with the mutation (Figure 1C). Because ψ-Ex contains a termination codon, the mRNA may be degraded by nonsense-mediated mRNA decay (NMD) as previously suggested (42, 43). To test this possibility, the cells were treated with puromycin to inhibit NMD (33). Inhibition of NMD was confirmed by the increase in abundance of a known NMD substrate (Supplementary Figure S1) (44, 45). Puromycin treatment resulted in an increase in mRNA with ψ-Ex to 38% of total *CFTR*, confirming that *CFTR* mRNA containing ψ-Ex is unstable and likely degraded by NMD (Figure 1C). Treatment of the cells with ASO-ψ, resulted in undetectable levels of ψ-Ex splicing (Figure 1C), demonstrating that ASO-ψ effectively blocks the use of the aberrant splice site, decreasing the amount of aberrantly spliced *CFTR* mRNA in a patient-derived lymphoblastoid cell line.

### ASO-ψ improves chloride secretion in human bronchial epithelial cells isolated from CF patients

We tested next whether ASO-ψ could improve chloride secretion, as defined by a forskolin-stimulated and bumetanide-inhibited I_eq_, in primary human bronchial epithelial (hBE) cells isolated from lung explants of CF patients (46–48). Cells derived from two patients compound heterozygous for *CFTR* c.3718-2477C>T and F508del-*CFTR* (donors 1 and 2) and one homozygous for the splice mutation (donor 3) were tested. Cells were differentiated on transwell filter plates and transfected with vehicle, ASO-C, or ASO-ψ. Four days later, equivalent currents (I_eq_) were measured using a TECC-24 workstation. Transepithelial voltage (Vt) and resistance (Rt) were recorded to calculate an I_eq_ attributable to forskolin-stimulated chloride secretion following the addition of benzamil to inhibit sodium currents mediated by epithelial sodium channels (ENaC) on the apical membrane (Figure 2A) (49–51). We also tested the activity of the potentiator VX-770, an FDA-approved therapeutic for patients with *CFTR* c.3718-2477C>T (27). Bumetanide, an inhibitor of the basolateral membrane chloride co-transporter NKCC1, was added 20 minutes after VX-770 to confirm that the measured current was chloride-specific (Figure 2A). All patient donor cells had similar chloride-specific currents following forskolin stimulation and treatment with ASO-C (plateau I_eq_=21; 18; 18 μA/cm^2^, donors 1-3 respectively) (Figure 2B, C left y-axis; Supplementary Table S2).

**Figure 2.**
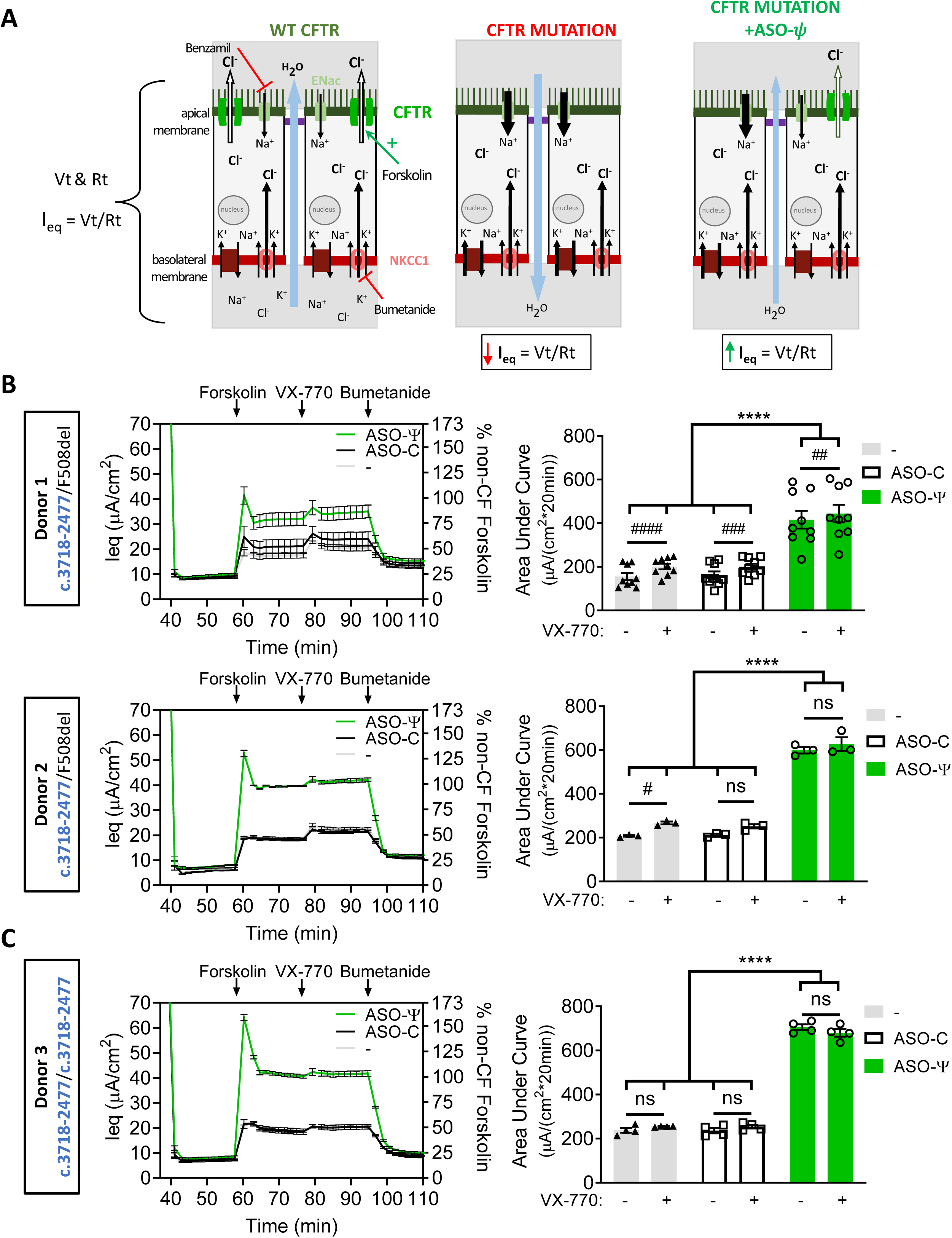
ASO-ψ treatment increases chloride-specific currents. (**A**) Diagram depicting chloride secretion in patient epithelial cells mediated by CFTR expression at the apical membrane. Equivalent current (I_eq_) is calculated by Vt and Rt measurements by electrodes in the apical and basolateral solutions. (left) Chloride secretion in cells with WT-CFTR. (middle) CFTR expression at the apical membrane is diminished by mutations such as *CFTR* c.3718-2477C>T, disrupting chloride secretion as indicated by a decrease in I_eq_. (right) Restored CFTR expression at the apical membrane mediated by ASO-ψ results in an increase in chloride secretion (increased I_eq)._ Analysis of differentiated hBE cells from **(B)** two patient donors with the *CFTR* c.3718-2477C>T mutation and F508del-*CFTR* (donor 1; donor 2) and (**C**) a patient donor homozygous for the *CFTR* c.3718-2477C>T mutation (donor 3). (Left) Average I_eq_ traces calculated from the TECC assay (left y-axis) and represented as a percent of the I_eq_ measured from non-CF cells following normalization to the average forskolin calculated currents (right y-axis) (average forskolin-specific I_eq_=41 μA/cm^2;^ donors 5, 6). Cells were treated with vehicle (grey line), ASO-C (black line), or ASO-ψ (green line). (Right) Quantification of the area under the curve (AUC) of cells treated with vehicle, ASO-C, or ASO-ψ following the addition of forskolin or forskolin + VX-770 (±SEM; N=9 (donor 1), N=3 (donor 2), N=4 (donor 3); two-way ANOVA; Tukey’s multiple comparisons test between treatment groups, ****p<.0001; Sidak’s multiple comparisons test within treatment groups, ^#^p<0.05, ^##^p<0.01, ^###^p<0.001, ^####^p<0.0001, ns=p>0.05).

Treatment of the cells with ASO-ψ resulted in a significant increase in the chloride current in all three donor cells relative to control treatment (plateau I_eq_=32 (p<0.05); 40 (p<0.0001); and 41 (p<0.0001) μA/cm^2^, donor 1-3 respectively) (Figure 2B, C left-y-axis, Supplementary Table S2). This activity, following treatment with ASO-ψ, reached levels similar to hBE cultures from non-CF donors (Figure 2B, C right y-axis; Supplementary Figure S2B, C; Supplementary Table S2). Overall, ASO-ψ treatment more than doubled chloride secretion relative to control ASO-C treatment in the compound heterozygotes as measured by the change in the area under the curve (ΔAUC) representing the 20 minute test period (ΔAUC I_eq_=255 μA/cm^2^_*_20min, donor 1; 386 μA/cm^2^_*_20min, donor 2; p<0.0001) and tripled chloride secretion in the homozygous patient cells (ΔAUC I_eq_=470 μA/20min*cm^2^, donor 3: p<0.0001) (Figure 2C; Supplementary Table S2). Comparatively, there was only a modest improvement in chloride secretion with VX-770 treatment and no apparent additive effect with ASO-ψ treatment in combination with C18.

The effect of ASO-ψ is specific to the *CFTR* c.3718-2477C>T splice mutation as ASO-ψ treatment of non-CF hBE cells and homozygous F508del-*CFTR* hBE cells had no effect on chloride secretion compared to the control (Supplementary Figure S2B, C; Supplementary Figure S3D, E). In contrast, treatment of F508del-*CFTR* cells with the corrector (C18) and potentiator (VX-770) resulted in a 3-fold increase in chloride secretion, similar to the increase seen with ASO-ψ treatment of *CFTR* c.3718-2477T hBE cells (Supplementary Figure S3D, E).

### ASO-ψ corrects splicing in *CFTR* c.3718-2477C>T patient-derived bronchial epithelial cells (hBEs)

To confirm that the improvement in chloride secretion following ASO-ψ treatment was due to an ASO-mediated block of the aberrant ψ-Ex splice site and correction of splicing to the native site, we assessed splicing in the cells by RT-PCR. In the homozygous donor cells, nearly 75% of *CFTR* mRNA contained ψ-Ex and treatment with ASO-ψ significantly reduced ψ-Ex inclusion to 11% of total *CFTR* mRNA (Figure 3A, B). This reduction in ψ-Ex splicing correlated with an increase in WT-*CFTR* RNA transcripts from *CFTR* c.3718-2477T to nearly non-CF (WT) levels (Figure 3C). Puromycin-treatment increased mRNA with the ψ-Ex exon and ASO-ψ significantly reduced this ψ-Ex splicing (Figure 3A, B). ASO-ψ treatment also resulted in an increase in the ratio of mature CFTR protein relative to total CFTR (Figure 3D, E) (52). ASO-mediated correction of splicing was dose-dependent with a half-maximal effective concentration (EC_50_) of 14.72 μM (Figure 3F, G). This dose-dependent effect on splicing correlated with an increase in chloride secretion (R^2^=0.7364; p=0.0031) (Figure 3H).

**Figure 3.**
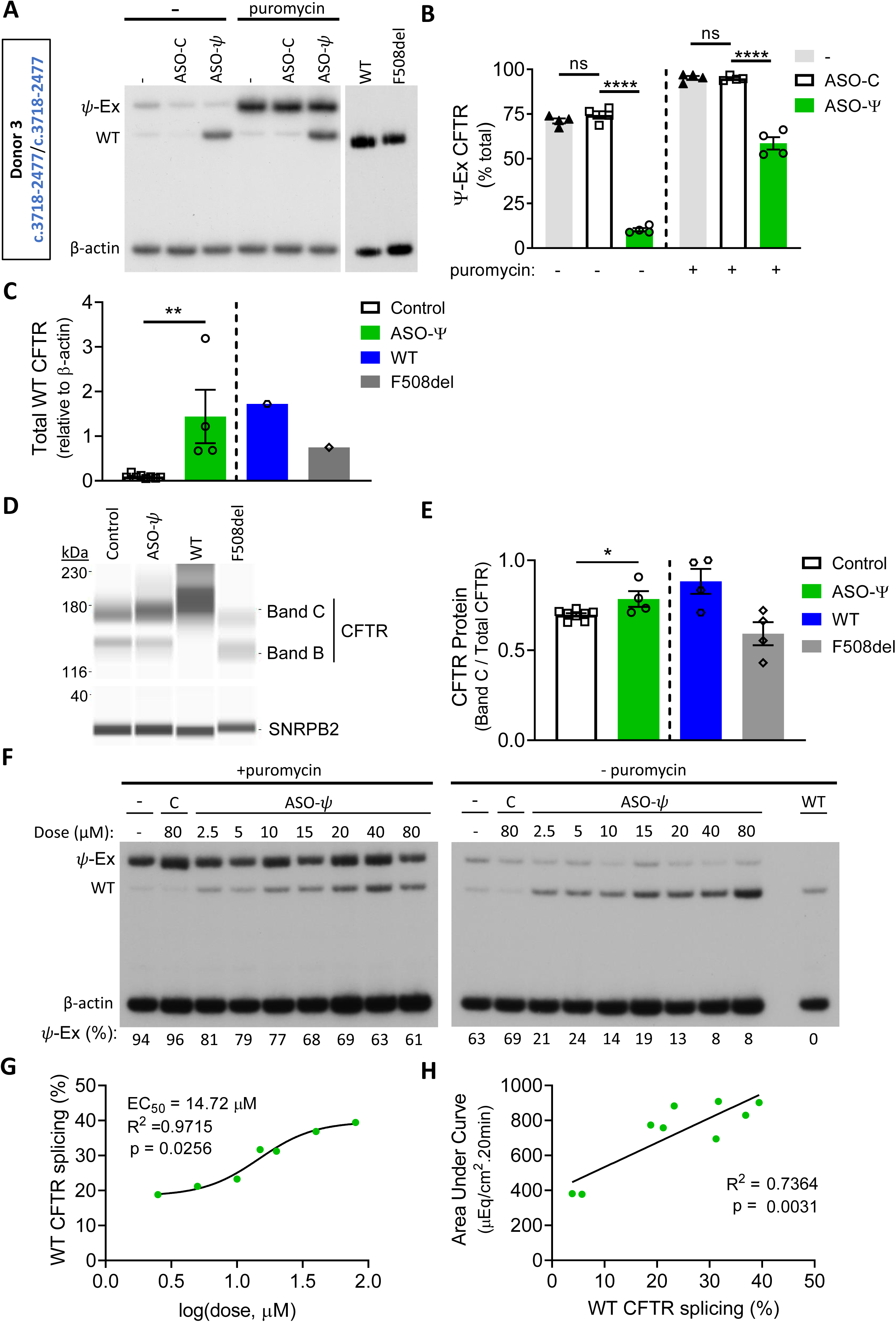
ASO-ψ treatment increases full-length *CFTR* mRNA abundance. **(A)** RT-PCR analysis of homozygous *CFTR* c.3718-2477C>T hBE cells (donor 3) treated with vehicle (-), ASO-C, or ASO-ψ and with or without puromycin. β-actin is shown as a control. **(B)** ψ-Ex splicing quantified as the percent of total *CFTR* (±SEM; N=4; Ordinary one-way ANOVA with Tukey’s multiple comparisons test, ****p<0.0001). **(C)** Total WT-*CFTR* from non-puromycin treated samples quantified relative to β-actin. (±SEM; N=8 control (ASO-C + (-)), N=4 ASO-ψ; Unpaired t-test, **p<0.01) **(D)** Immunoblot analysis of CFTR protein isolated from homozygous patient cells treated with ASO-ψ or control (vehicle + ASO-C). SNRPB2 is a loading control. **(E)** Quantification of the B + C / Total CFTR from (C). (±SEM; N=8 control (ASO-C + (-)), N=4 ASO-ψ; Unpaired t-test, *p<0.05) **(F)** RT-PCR analysis of *ψ*-Ex splicing in *CFTR* c.3718-2477C>T homozygous cells treated with ASO-ψ or ASO-C at concentrations indicated. β-actin represents a control for RNA expression. Aberrantly spliced RNA was quantified as percent *ψ*-Ex splicing and is shown below each lane. **(G)** WT-*CFTR* splicing in the cells treated with puromycin, from (F), related to ASO-ψ dose. The half-maximal effective concentration (EC_50_) was calculated after fitting the data using the normalized response and variable slope. **(H)** The calculated area under the curve after forskolin treatment correlated to WT-*CFTR* splicing, shown in (E) in cells treated with puromycin (simple linear regression).

In the compound heterozygous donor cells, a significant decrease in ψ-Ex splicing was observed, though ψ-Ex mRNA could only be detected following puromycin treatment to block NMD (Supplementary Figure S4A, B). In these cells, the majority of the WT products likely derive from the F508del-*CFTR* allele. Thus, to confirm the effect of ASO-ψ in these cells, we quantified *CFTR* mRNA derived solely from the *CFTR* c.3718-2477T allele by real-time PCR using a probe that does not bind to F508del-*CFTR* mRNA. We observed a significant increase in total *CFTR* c.3718-2477T mRNA as predicted with an increase in the abundance of the stable, full-length *CFTR* mRNA and a decrease in the unstable *CFTR* containing ψ-Ex (Supplementary Figure S4C). No change in mRNA from the F508del-*CFTR* allele was observed after ASO treatment indicating that the increase in correct splicing mediated by ASO-ψ was specific to *CFTR* c.3718-2477T (Supplementary Figure S4D).

### ASO-induced correction of splicing is long-lasting in cultured hBE cells

To assess the longevity of ASO-ψ activity we treated patient-derived hBE cells (homozygous *CFTR* c.3718-2477T donor 3) with ASO-ψ or ASO-C and measured chloride currents and splicing 4, 8, and 16 days later. ASO-ψ treatment resulted in a significant and sustained increase in chloride currents compared to cells treated with ASO-C for the same treatment period with a modest increase in activity at 8 and 16 days (p=0.0023, p=0.0029) (Figure 4A, B; Supplementary Table S3). After 16 days of ASO treatment, baseline, post-benzamil currents significantly increased to a level comparable to that found in non-CF hBEs, which could be further indicative of a corrective change in transport phenotype to normal levels (Figure 4A, Supplementary Table S2, S3). The enduring secretory currents correlated with a similarly prolonged correction of splicing (Figure 4C) and elevation of wild-type spliced *CFTR* mRNA (Figure 4D). Taken together these results show that ASO-ψ has a long-lasting effect in patient cells, with sustained correction of aberrant splicing and rescue of CFTR function.

**Figure 4.**
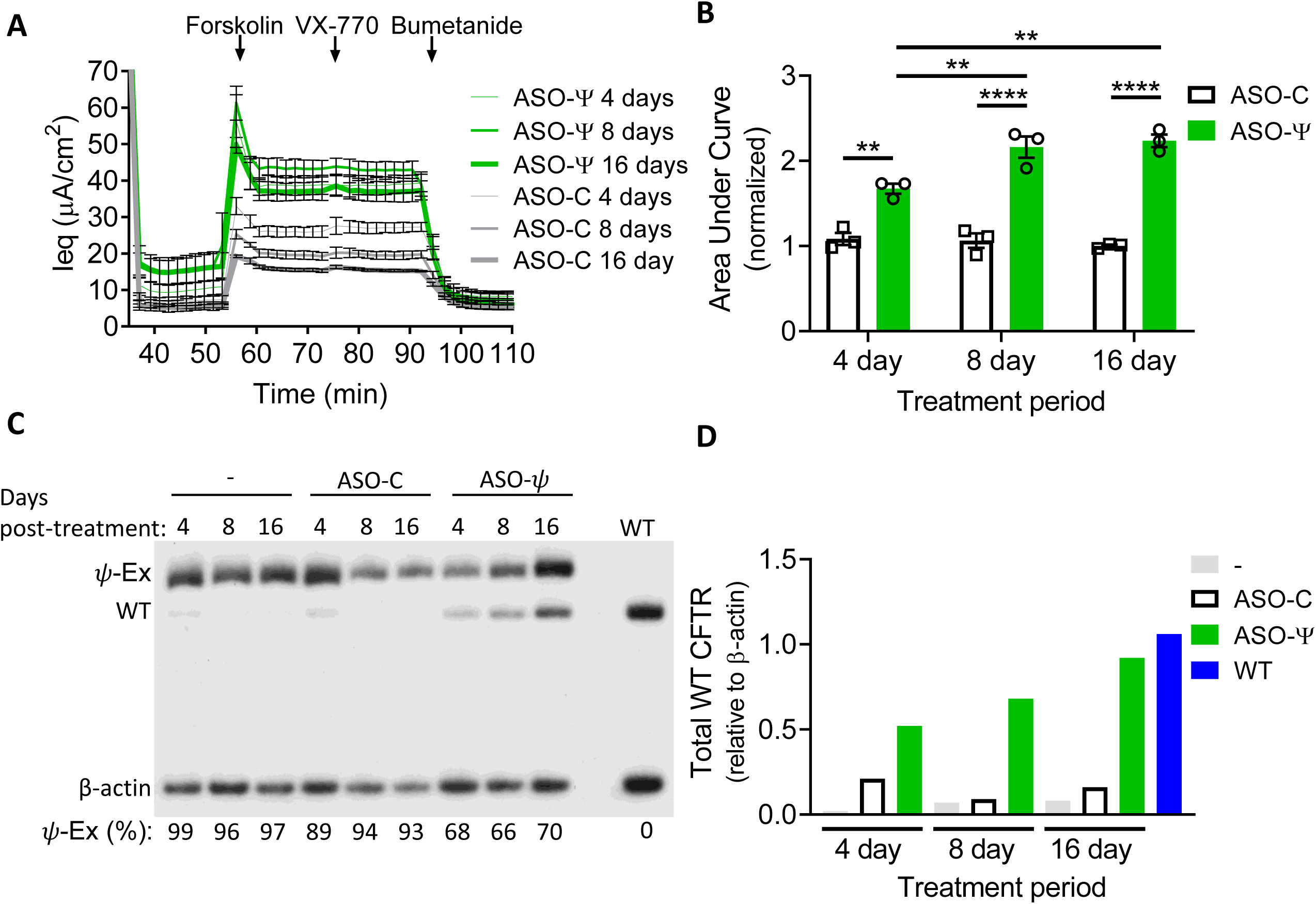
ASO-ψ rescues chloride current and splicing for 16 days. **(A)** Average I_eq_ traces of hBE cells from a homozygous *CFTR* c.3718-2477C>T patient (donor 3) were treated with ASO-ψ or ASO-C for 4, 8, and 16 days and analyzed for chloride secretion. **(B)** The area under the curve was calculated after forskolin addition and data was normalized to vehicle-treated cells (±SEM; N=3; two-way ANOVA with Tukey’s multiple comparisons test, **p<0.01, ****p<0.0001). **(C)** RT-PCR analysis of *CFTR* splicing from RNA isolated from the cells after functional analysis and puromycin treatment. β-actin is a control for total RNA level. RNA was quantified as percent *ψ*-Ex splicing and shown below each lane. **(D)** Quantification of WT-*CFTR* relative to β-actin from panel (C).

### ASO-ψ treatment increases apical membrane conductance

Chloride transport into and out of epithelial cells relies on channels, transporters, and pumps in both the apical and basolateral membranes (2). In airway cells, chloride entry into the cell through the basolateral membrane co-transporter NKCC1 is coupled to secretion across the apical membrane via CFTR to sustain ion and fluid transport (Figure 2A) (2, 53, 54). Historically, transepithelial impedance analysis has been used to investigate the relative contribution channels in the two membranes have in chloride transport (40, 41, 51). To confirm that ASO-ψ specifically recovers CFTR channel activity at the apical membrane, we performed transepithelial impedance analysis on homozygous patient hBE cells following treatment with ASO-ψ. ASO-ψ nearly doubled apical conductance compared to ASO-C treatment after stimulation with forskolin (p<0.0001) (Figure 5B; Supplementary Figure S5). Treatment with VX-770 with or without ASO-ψ did not result in a significant increase in apical conductance (Figure 5B). The ASO-ψ-mediated increase in apical conductance was dependent on chloride, as the increase in conductance was reversed by the addition of an inhibitor of chloride conductance, GlyH-101 (Figure 5B) (55, 56). The correlation between an increase in apical conductance and treatment with the *CFTR*-specific ASO-ψ is strong evidence the drug is rescuing CFTR expression.

**Figure 5.**
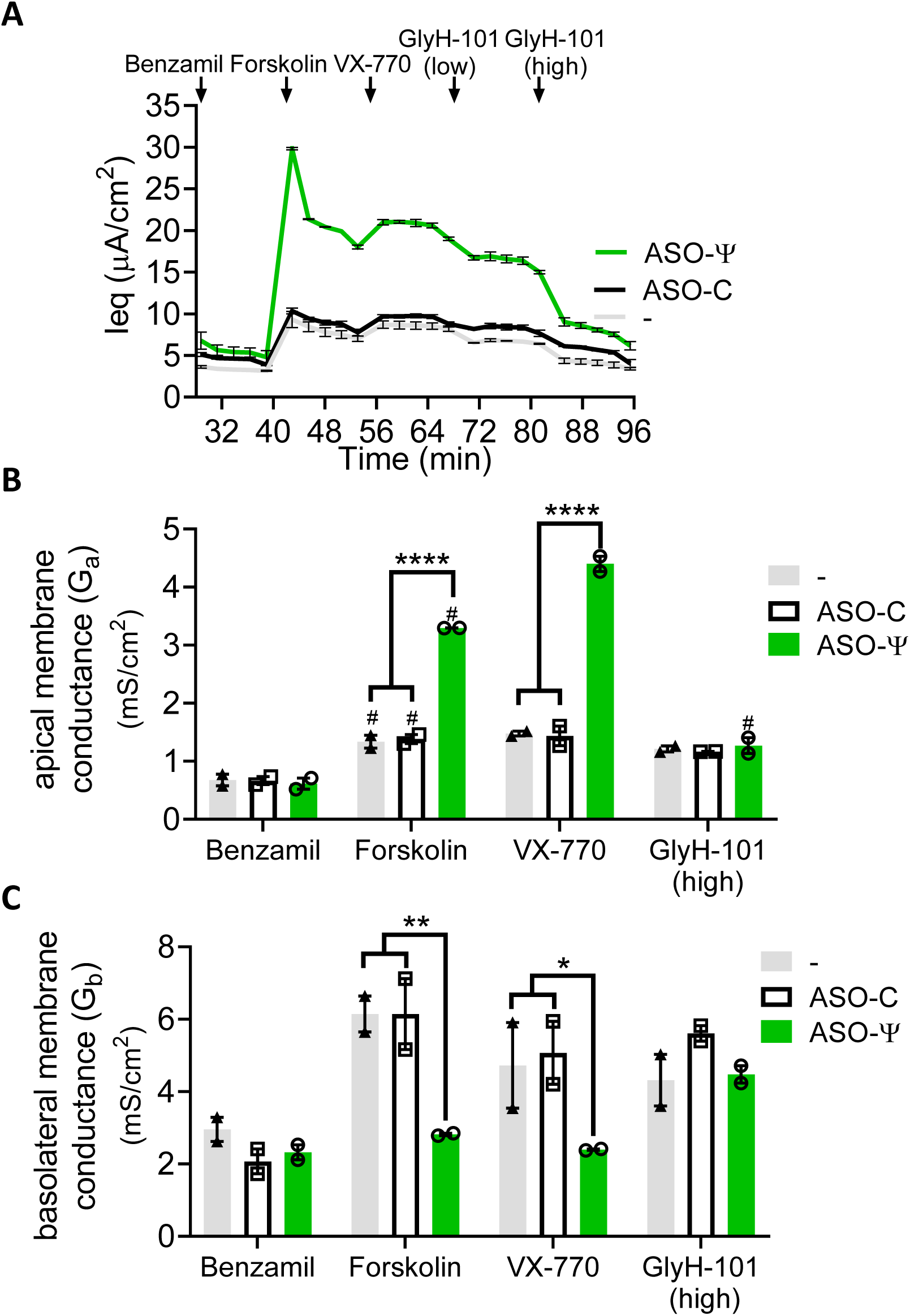
ASO-ψ increases apical membrane conductance. (**A**) Average I_e_q traces were calculated from hBE cells isolated from a CF patient donor homozygous for *CFTR* c.3718-2477C>T (donor 3) treated with vehicle (grey line), ASO-C (black line), or ASO-ψ (green line). Average (**B**) apical conductance (G_a_) and (**C**) basolateral conductance (G_b_) were calculated by fitting impedance spectra data from the patient epithelial cells to equivalent circuit equations that model epithelial membranes after benzamil, forskolin, VX-770, and GlyH-101 additions. The four data points following the initial compound addition were averaged (±SEM, N=2; two-way ANOVA; Tukey’s multiple comparisons test between treatment groups, *p<0.05, **p<0.01, ****p<0.0001; Sidak’s multiple comparisons test within treatment groups, #p<0.05).

Treatment with ASO-ψ also resulted in a greater than 2-fold reduction in basolateral membrane conductance after stimulation with forskolin compared to ASO-C-treated cells (p=0.005) (Figure 5C). Basolateral Na/K ATPase activity has been reported to be two-fold higher in CF patients (57, 58), which is consistent with the elevated conductance we observed in control-treated cells (Figure 5C). Although more testing is needed to assess Na/K ATPase directly, the normalization of basolateral conductance with ASO-ψ treatment is consistent with a correction of the ion secretion pathway disrupted by the loss of CFTR. Together, these results provide evidence that the ASO-ψ-dependent increase in chloride secretion is mediated by elevated CFTR activity.

### ASO-ψ is superior to other CFTR modulators at improving chloride secretion in *CFTR* c.3718-2477C>T patient cells

Potentiator and corrector therapeutics are available to patients with the *CFTR* c.3718-2477C>T mutation (27, 28). An ASO therapeutic likely would be used in place of or in combination with these drugs. Thus, we compared the effect of ASO-ψ and two of the current CFTR drug treatments, VX-770 (ivacaftor) and C18, an analog of the corrector VX-809 (lumacaftor) on chloride transport. Rescue of chloride current was analyzed as before with the exception that three days after ASO-transfection (24 hours before functional analysis) cells were treated with C18 or vehicle.

Modulator treatment (C18+VX-770) resulted in a 2.1-fold increase in chloride secretion in the *CFTR* compound heterozygous cells (Figure 6A) (donor 1, p<0.0001; donor 2, p<0.0001), presumably due to targeting of the F508del-*CFTR* protein. In contrast, C18+VX-770 treatment had no effect on homozygous *CFTR* c.3718-2477C>T donor cells (p=0.64) (Figure 6 right). This result suggests that the modulator treatment does not act on WT-*CFTR* produced by residual correct splicing of *CFTR* c.3718-2477T RNA nor on any truncated protein produced from mRNA with ψ-Ex.

**Figure 6.**
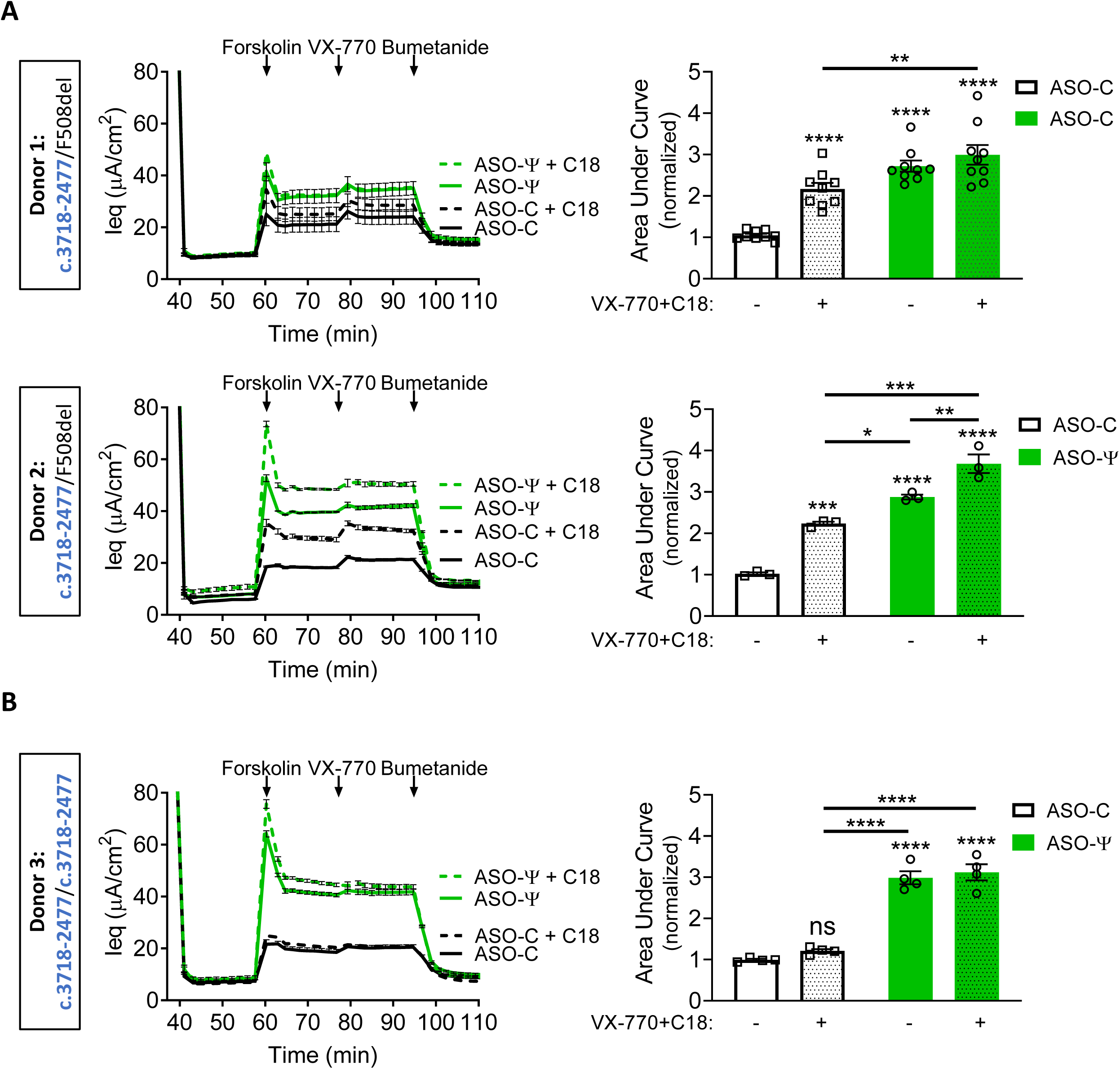
ASO-ψ increases chloride secretion above that achieved with current CFTR modulators. (left) Average I_eq_ traces of hBE cells isolated from (**A**) patients heterozygous for *CFTR* c.3718-2477C>T and F508del-*CFTR* (donor 1 and donor 2) and (**B**) a patient homozygous for *CFTR* c.3718-2477C>T (donor 3) treated with DMSO (.01%) (solid lines) or C18 (6μM) (hashed lines). Cells were treated with ASO-C (black lines), or ASO-ψ (green lines). (Right) Quantification of the area under the curve of cells treated with (stippled) or without (non-stippled) CFTR modulators (C18/VX-770) with ASO-C (white) or ASO-ψ (green). Data was normalized to the AUC of forskolin treatment in cells treated with vehicle alone (±SEM; N=9 (donor 1), N=3 (donor 2), N=4 (donor 3); two-way ANOVA with Tukey’s multiple comparisons test, *p<0.05, **p<0.001, ***p<0.001, ****p<0.0001, ns=p>0.01).

ASO-ψ treatment resulted in a 1.3-fold increase in chloride channel activity compared to the VX-770+C18 treatment in cells from the compound heterozygote donor 2 (p=0.0232), whereas treatment of the homozygous *CFTR* c.3718-2477C>T donor with ASO-ψ resulted in a 2.5-fold increase compared to modulator treatment (p<.0001). Finally, combining ASO-ψ with the modulators showed significantly greater recovery of activity relative to modulator alone for all three donors (donor 1, 1.4-fold, p=0.0041; donor 2, 1.6-fold, p<0.0001; homozygous donor 3, 2.6-fold, p<0.0001). However, the triple combination treatment only provided a significant increase in activity over ASO-ψ-treatment alone in donor 2 (p=0.0066, 1.3-fold) (Figure 6A).

Importantly, the rescue of channel activity achieved by the triple combination of VX-770, C18 and ASO-ψ in all donors (donor 1, 2.9-fold, donor 2, 3.6-fold, donor 3, 3.1-fold; p<0.0001) (Figure 5) was comparable to the level achieved by modulator treatment in hBE cells isolated from patients homozygous for the F508del-*CFTR* mutation (donor 4, 3.4-fold, p<0.0001) (Supplementary Figure S3D, E), providing further support for the potential efficacy of ASO-ψ in patients. As expected, ASO-ψ treatment had no effect on the F508del-*CFTR* cells (p=0.95, p=0.2) (Supplementary Figure S3D, E). Together these results demonstrate that, in our assay, ASO-ψ improves chloride secretion in patient-derived *CFTR* c.3718-2477C>T hBE cells to a greater degree than VX-770 alone (Figure 2) or in combination with C18 (Figure 6) and that, in some cases, modulator drugs and ASOs could have a partially additive therapeutic benefit.

## DISCUSSION

The *CFTR* c.3718-2477T splicing mutation is one of the more common CF-causing variants, yet none of the currently available treatments for CF target the splicing defect directly. A previous study reported on ASO-mediated inhibition of *CFTR* c.3718-2477T aberrant splicing in a CFTR expressing mini-gene system with the mutation, though no further work on this mutation has been done to show that ASOs can be used to recover CFTR activity in epithelial cells (25). Here, we demonstrate that ASO-ψ, an ASO designed to block splicing to the aberrant ψ-Ex splice site created by the *CFTR* c.3718-2477C>T mutation, redirects splicing to the correct, wild-type splice site, increases forskolin-stimulated apical membrane conductance and restores chloride secretion in CF patient-derived bronchial epithelial cells. This restoration of chloride secretion is comparable to that achieved in F508del-*CFTR* homozygous patient cells treated with the potentiator CF drug VX-770 and C18 (an analog of the corrector CF drug VX-809), a drug combination mimicking the clinically-approved treatment (VX-770+VX-809) currently used to treat CF patients with the F508del mutation. This result suggests that ASO-ψ could have similar efficacy in CF patients with the *CFTR* c.3718-2477T mutation. Comparatively, treatment of homozygous *CFTR* c.3718-2477T cells with VX-770 alone, which is approved for CF patients with several different splice mutations, did not achieve significant rescue of chloride secretion, suggesting that ASO-ψ could offer an improvement in the standard of care for these patients (27). We also show that ASO-ψ works together with approved corrector and potentiator drugs to increase channel activity in cells from patients that are compound heterozygous for the splice mutation and the common F508del-*CFTR* mutation, thus broadening the treatment options and improving efficacy for CF patients with the *CFTR* c.3718-2477C>T mutation.

Antisense technology is being explored as a therapeutic for CF, including strategies to elevate CFTR expression (59, 60), modulate other channels in the ion secretion pathway (61, 62), and target F508del-CFTR (63). Although splicing mutations account for a significant portion of CF cases, splice-switching ASOs have only been tested for two such mutations, *CFTR* c.3718-2477 C>T and c.2647+5G>A (11, 25). Igreja et al. (11), similar to previous work on *CFTR* c.3718-2477T-targeting ASOs mentioned above (25), used a *CFTR* cDNA mini-gene expression system in HEK-293 cells to demonstrate an ASO-mediated correction of *CFTR* aberrant splicing caused by the *CFTR* c.2647+5G>A mutation. They went further to demonstrate that the ASO-corrected splicing resulted in an increase in CFTR activity in an iodide efflux assay (11). Our study broadly advances the potential of ASOs for CF splicing mutations by showing successful recovery of endogenous *CFTR* splicing, protein expression, and chloride secretion in patient-derived hBE cells with the *CFTR* c.2718-2477C>T mutation. Further, we show that ASO-mediated correction recovers CFTR function to levels comparable to current approved CF modulators demonstrating the efficacy of ASO therapeutics.

*CFTR* c.3718-2477C>T, categorized as a class V *CFTR* mutation, produces some functional CFTR as a result of the use of the native exon 22 5’ donor splice site (64). Patients with the c.3718-2477C>T splice mutation have a spectrum of disease severity, often presenting with a less severe form of the disease (16–18, 65, 66). Pulmonary disease severity has been linked to the amount of aberrant splicing in patients, with small differences in ψ-Ex splicing correlating with comparatively large differences in pulmonary function, as determined by the percent predicted forced expiratory volume (ppFEV_1_) (15, 67). Based on these previous studies, the increase in correct splicing that we observed following ASO-ψ treatment (Figure 3) has the potential to result in a clinically relevant recovery of function (15, 67). In hBE cell cultures, we find that ASO-ψ treatment reached levels seen in non-CF patient cells (Figure 2B, C right y-axis; Supplementary Figure S2). However, this comparison must be considered with the caveat that there are other phenotypic differences between the CF and non-CF hBE cells such as differences in baseline transport in pre and post-benzamil currents (Supplementary Figure S2A). Thus, we consider our comparison of recovery to current CF modulators within the same donor cells to be a better predictor of therapeutic efficacy (68–71).

Because the *CFTR* c.3718-2477C>T allele yields some correctly spliced mRNA and thereby wild-type, full-length CFTR, the potentiator molecule VX-770 has been approved for the treatment of patients with the mutation (27). VX-770 has been reported to increase the activity of some CFTR mutants and wild type CFTR channels, although its effect on WT-*CFTR* is variable (72, 73). In our study VX-770 treatment did not elevate chloride current in non-CF hBE cells (Supplementary Figure S2B, C) and provided little improvement in channel activity in c.3718-2477T cells (Figure 2), although it is possible that a VX-770 effect could be obscured by hyperphosphorylation of CFTR (74, 75). Likewise, data from clinical trials showed only modest improvement in endpoints for patients with the splice mutation (29, kalydecohcp.com). Thus, our ASO treatment offers a potential advantage over this currently available treatment as evidenced by the ∼3-fold increase in chloride current in patient cells, a significant improvement compared to VX-770 alone (Figs 2,5). For compound heterozygous patients, who have the splice mutation in combination with the F508del-*CFTR* mutation, multiple corrector compounds (e.g. VX-809/lumacaftor) that specifically target the folding and trafficking defects produced by the F508del-*CFTR* mutation have been approved for use in combination with potentiators (9, 10, 38, 47, 77). In our study, a combination treatment of compound heterozygous c.3718-2477T/F508del patient cells with a VX-809 analog, C18, and VX-770 resulted in a 2-fold increase in CFTR channel currents (Figure 6). Comparatively, ASO-ψ treatment resulted in a 3-fold increase in chloride secretion in the same patient cells, a significant improvement in correction over the approved drugs in one of the patient cell models.

Importantly, treatment with ASO-ψ in combination with the modulators VX-770 and C18 increased chloride secretion up to 3.5-fold, a significantly greater improvement than modulator treatment alone, supporting the use of an ASO treatment in combination with therapies available for patients with the splice mutation and another *CFTR* mutation that would respond to corrector and potentiator therapies (Figure 6). Interestingly, the treatment of hBE cells with ASO-ψ and modulators was not fully additive in the compound heterozygous donors. It is possible that the maximal effect is limited by intracellular regulation of the chloride secretion process. Further investigation is needed to understand this effect (69). Nevertheless, this level of functional recovery, achieved with ASO alone in patients homozygous for the splice mutation and in combination with modulators in the compound heterozygotes, is comparable to levels achieved by VX-770 with C18 treatment in F508del-*CFTR* homozygous patient cells (Supplementary Figure S3E). This result suggests that ASO-ψ could be just as effective a drug for *CFTR* c.3718-2477C>T patients as the clinically successful corrector/potentiator drugs are for F508del-*CFTR* CF patients. The newest correctors clinically available to CF patients, tezacaftor (VX-661) and elexacaftor (VX-445), have shown improved efficacy compared to lumacaftor (VX-809) (9, 10). Whether ASO treatment would result in a significant increase in CFTR channel activity compared to these newer treatments remain to be determined. However, the semi-additive therapeutic effect of ASO-ψ and C18 observed in this study provides evidence that ASO treatment could have an additional benefit in combination with any corrector compound.

ASOs have been shown to have long-lasting effects *in vivo*, with treatment durations lasting weeks to months before further dosing in patients (20, 21, 78, 79). Similarly, we observe that one dose of ASO-ψ resulted in a sustained rescue of *CFTR* splicing and function for up to 16 days in hBE cells, demonstrating the potential long-lasting effect of ASOs in human epithelial cells (Figure 4). If ASOs have a similar lasting effect *in vivo*, the frequency of dosing may be an advantage to this therapeutic approach. By comparison, current CFTR modulators have daily prescribed dosing (80, 81).

In order to be considered for therapy, an ASO drug must be effectively delivered to clinically relevant cells. The airway epithelial cells are one of the primary targets for CF therapeutics. The clinical potential of ASOs delivered directly to the respiratory system has been demonstrated for asthma and other inflammatory lung conditions (82–85). Unlike *in vitro* cell culture systems, which typically require transfection reagents for efficient delivery, naked ASOs have been successfully delivered to the lung, where they access multiple cell types including airway epithelial cells (62, 83, 86, 87). Specific to CF, aerosolization of ASOs has shown promise as an effective delivery modality to CFTR expressing lung epithelial cells in both a CF-like lung disease model in mice as well as CF patients (86, 88, 89). In recent clinical trials, treatment with an ASO drug candidate, eluforsen, which specifically targets the F508del-*CFTR* mutation, resulted in improved chloride and sodium transport, as measured by a nasal potential difference (NPD), in CF patients when delivered via bilateral intranasal administration three times a week for four weeks (63). These studies provide evidence of the delivery, efficacy, and safety of ASOs in targeting *CFTR* when delivered to the lung in both mouse models of CF and CF patients. Additionally, although patients with *CFTR* c.3718-2477 C>T mutations have mild phenotypes outside of lung disease, systemic delivery of ASOs may also be considered to correct CFTR expression in additional tissue types. Intravenous and subcutaneous injections are currently used to treat patients with ASO therapeutics and a bioavailable ASO formulation has been reported to target the gut epithelia, a tissue with relevance for CF patients (90–92).

In summary, we show that ASO treatment directed to the *CFTR* c.3718-2477C>T splice mutation blocks splicing to the cryptic splice site resulting in an increase in correctly spliced *CFTR* mRNA. The ASO increases correct splicing to a level predicted to be clinically relevant in CF patients based on previous reports correlating an increased level of aberrant splicing with decreased patient lung function (15, 67). This conclusion is supported by our results showing a 3-fold increase in chloride current after treatment with ASO-ψ. This correction is significantly above that achieved in comparison to current modulators approved for patients with the *CFTR* c.3718-2477C>T splice mutation. Our results also provide evidence that ASO treatment could provide an additional treatment option in combination therapies currently being tested for compound heterozygous CF patients with the splice mutation.

## Supplementary Data

Supplementary Data are available online.

## Acknowledgments

The authors thank Dr. Scott Randell, University of North Carolina School of Medicine, for providing the patient cells and Dr. Adrian Krainer, Cold Spring Harbor Laboratory, for the 4G3 antibody. We are grateful to Drs. Craig Hodges and Anne Harris, Case Western Reserve University, for their intellectual contribution and Rusudan Kotaria, Hui Li and Matt Green, for technical support.

## Author contributions

M.L.H. and R.J.B. initiated and supervised the study; W.E.M, M.L.H, and R.J.B. designed the experiments; W.E.M. performed all the experiments; W.E.M., M.L.H., and R.J.B. analyzed the data; W.E.M and M.L.H wrote the manuscript with contributions from all authors.

