## Supplementary Figure S1 for "Antisense oligonucleotide-mediated correction of *CFTR* splicing improves chloride secretion in cystic fibrosis patient-derived bronchial epithelial cells"

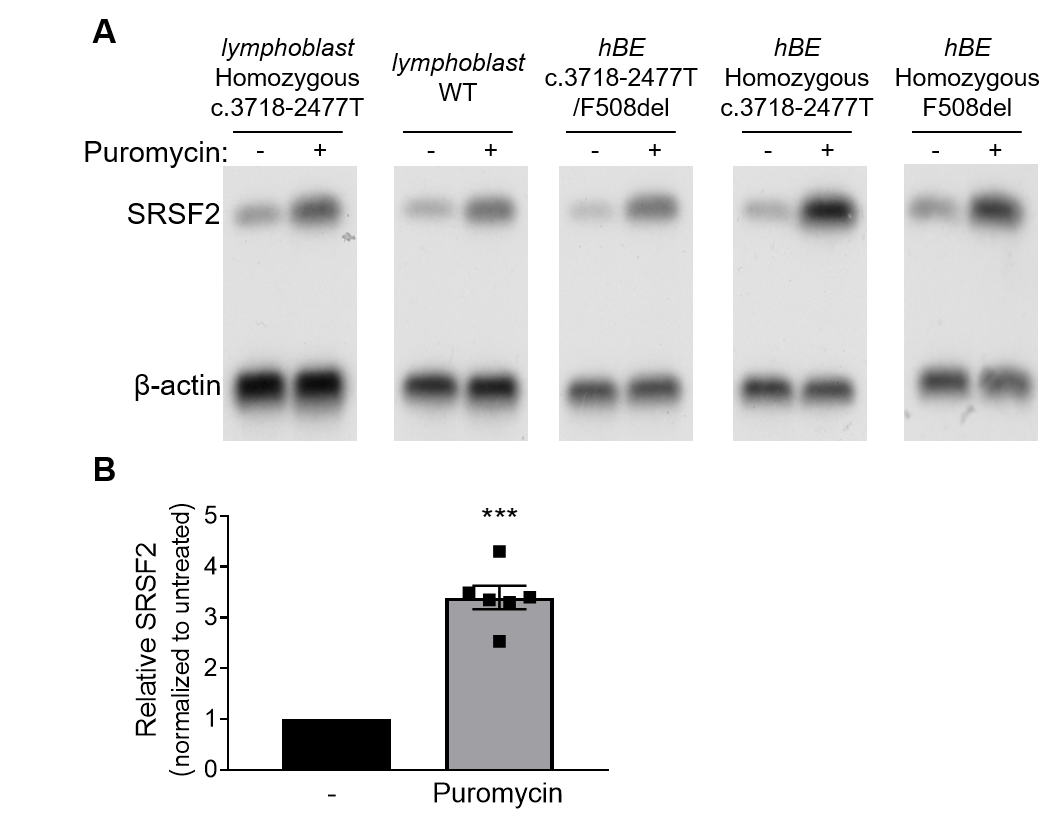

**Supplementary Figure S1**. Puromycin treatment of lymphoblasts inhibits nonsense-mediated mRNA decay of SRSF2 mRNA isoform with premature termination codon. **(A)** An RT-PCR analysis of SRSF2, a known substrate of NMD, expression in CFTR expressing cell models used in the study treated (+) or untreated (-) with puromycin (200μg/ul) for 9 hours (lymphoblasts) or 6 hours (hBEs). β-actin was used as a control for RNA expression. **(B)** SRSF2 expression after puromycin treatment was quantified and normalized to the untreated samples (±SEM; N=5; one sample t-test, ***p<0.001).
