## Supplementary Figure S2 for "Antisense oligonucleotide-mediated correction of *CFTR* splicing improves chloride secretion in cystic fibrosis patient-derived bronchial epithelial cells"

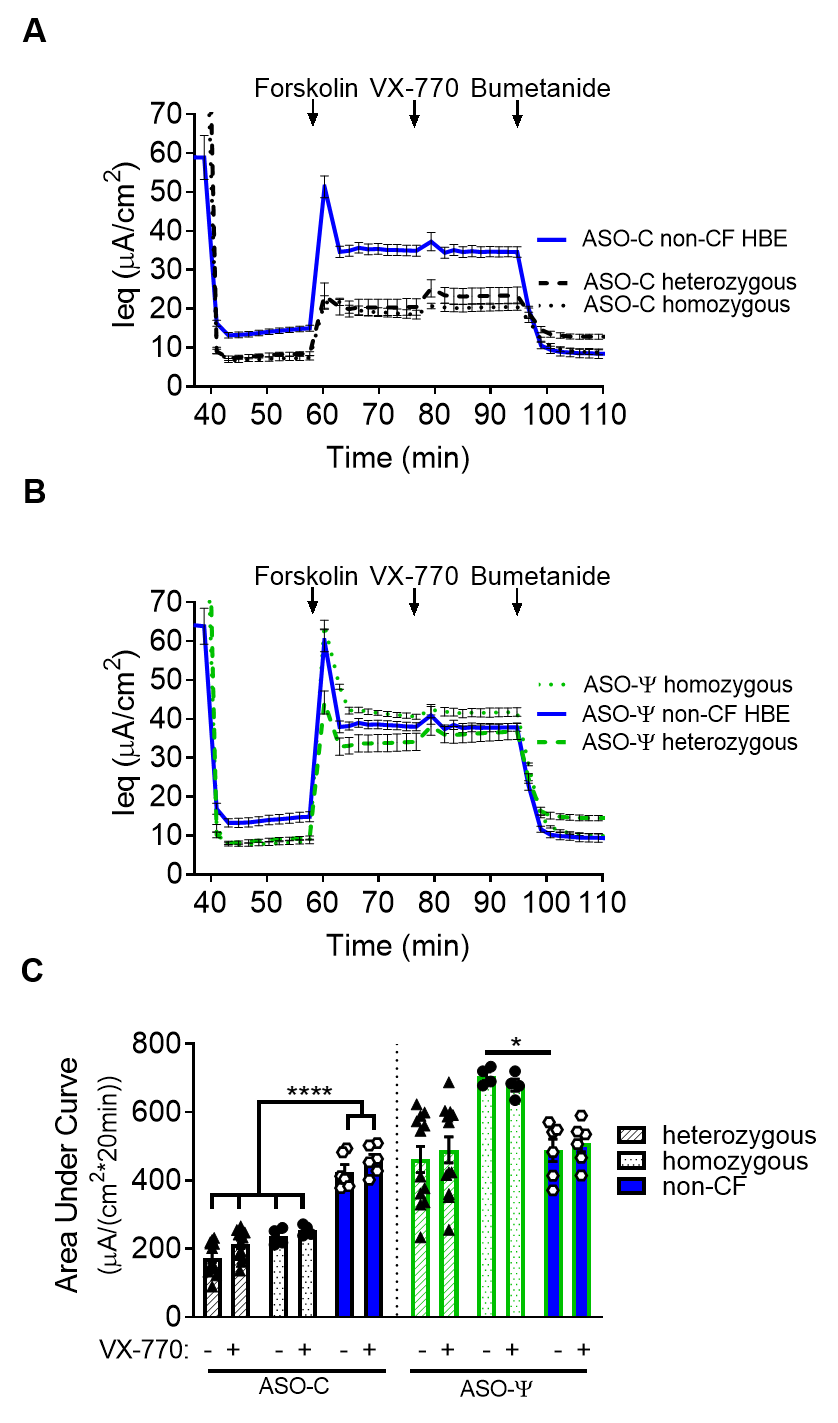

**Supplementary Figure S2.** Chloride currents in hBE cells from *CFTR* c.3718-2477T donor and non-CF donor cells. **(A)** Average I_eq_ traces from hBE cells from non-CF donors (donor 5 and donor 6) (blue line), compound heterozygous donors 1 and 2 and a homozygous donor 3 cells treated ASO-C (black lines) and **(B)** ASO-ψ. **(C)** Area under the curve of the forskolin treatment period was quantified for each genotype (±SEM; heterozygous N=12, homozygous N=4, non-CF N=6; Ordinary one-way ANOVA, Tukey’s multiple comparisons test, *p<.05, ****p<0.0001).
