## Supplementary Figure S3 for "Antisense oligonucleotide-mediated correction of *CFTR* splicing improves chloride secretion in cystic fibrosis patient-derived bronchial epithelial cells"

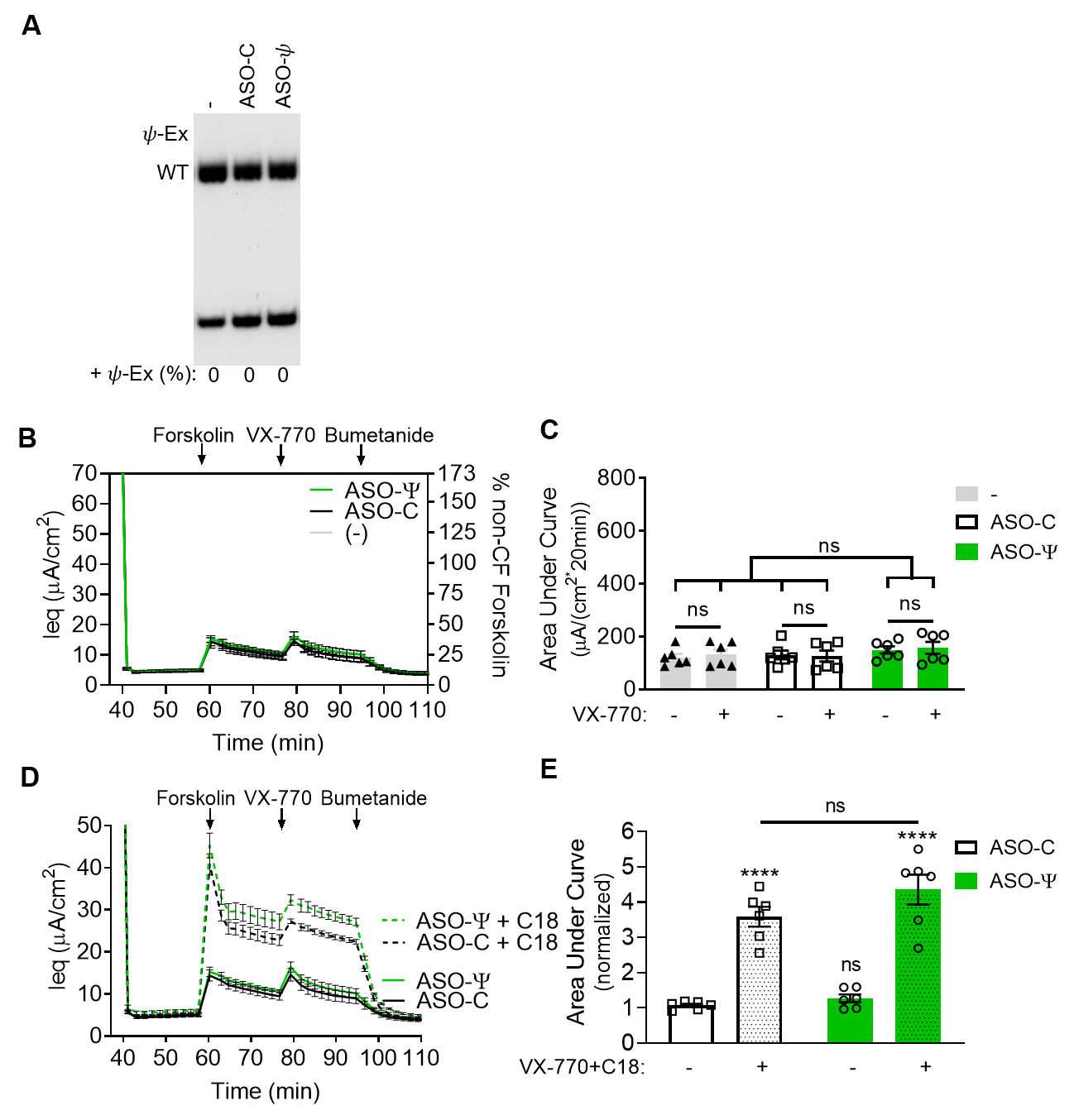

**Supplementary Figure S3.** Corrector (C18) and potentiator (VX-770), but not ASO-ψ treatment improves chloride channel activity in F508del-*CFTR* cells. **(A)** RT-PCR analysis of homozygous F508del-*CFTR* patient hBE cells (donor 4) treated with vehicle, ASO-C, or ASO-ψ. β-actin was included as a control for total RNA level. Splicing was quantified as the percent of RNA transcripts with the pseudo-exon insert [+ψ-Ex/((+ψ-Ex)+WT)x100)] and is shown below each lane. **(B)** Average I_eq_ traces calculated from the TECC assay (left y-axis) or normalized to percent non-CF average forskolin-specific currents (right y-axis) (average forskolin-specific I_eq_=41 μA/cm^2;^ donors 5, 6) from F508del-*CFTR* homozygous hBE cells treated with vehicle (grey lines), ASO-C (black lines), or ASO-ψ (green lines). **(C)** Area under the curve of the forskolin (unhashed) or forskolin + VX-770 (hashed) treatment period was quantified for samples treated with vehicle, ASO-C, or ASO-ψ (±SEM; N=6; Ordinary one-way ANOVA; Tukey’s multiple comparisons test between groups; Sidak’s multiple comparisons test within groups, ns=p>0.01). **(D)** Average I_eq_ traces from F508del-*CFTR* homozygous hBE cells treated with ASOs as before and with DMSO (solid lines) or C18 (6 μM, dashed lines). **(E)** Area under the curve analysis of cells treated with (hashed) or without (unhashed) CFTR modulators (C18, VX-770) along with ASO-C (white) or ASO-ψ (green) was quantified. Data was normalized to the quantification for forskolin treatment in vehicle treated samples (±SEM; N=6; Ordinary one-way ANOVA, Tukey’s multiple comparisons test, ****p<0.0001, ns=p>0.01).
