## Supplementary Figure S4 for "Antisense oligonucleotide-mediated correction of *CFTR* splicing improves chloride secretion in cystic fibrosis patient-derived bronchial epithelial cells"

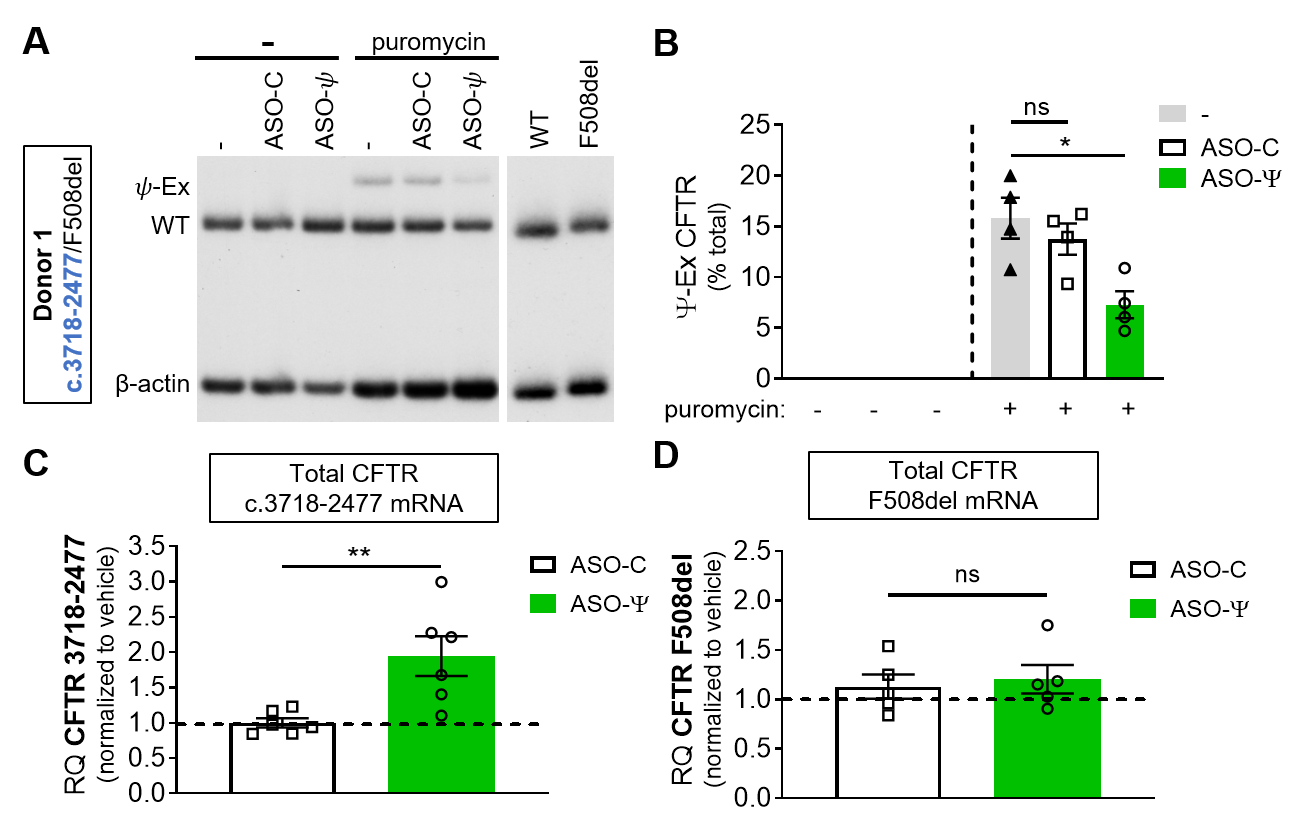

**Supplementary Figure S4**. ASO-ψ treatment increases full-length *CFTR* mRNA in compound heterozygous patient cells**.** (**A**) RT-PCR analysis of compound heterozygous hBE cells (c.3718-2477 T/F508del) treated as in Fig 3A. (**B**) ψ-Ex splicing quantified as the percent of total *CFTR*. (±SEM; N=4; Ordinary one-way ANOVA with Tukey’s multiple comparisons test, *p<0.05) (**C**) RT-qPCR analyses of total *CFTR* mRNA from the *CFTR* c.3718-2477C>T allele. (±SEM; N=6; t-test, **p<0.01) (**D**) F508del-*CFTR* isolated from the compound heterozygous patient (donor 1) treated as above (±SEM; N=5; t-test, ns=p>0.05). Relative quantity (RQ) was normalized to vehicle treatment.
