## Supplementary Figure S5 for "Antisense oligonucleotide-mediated correction of *CFTR* splicing improves chloride secretion in cystic fibrosis patient-derived bronchial epithelial cells"

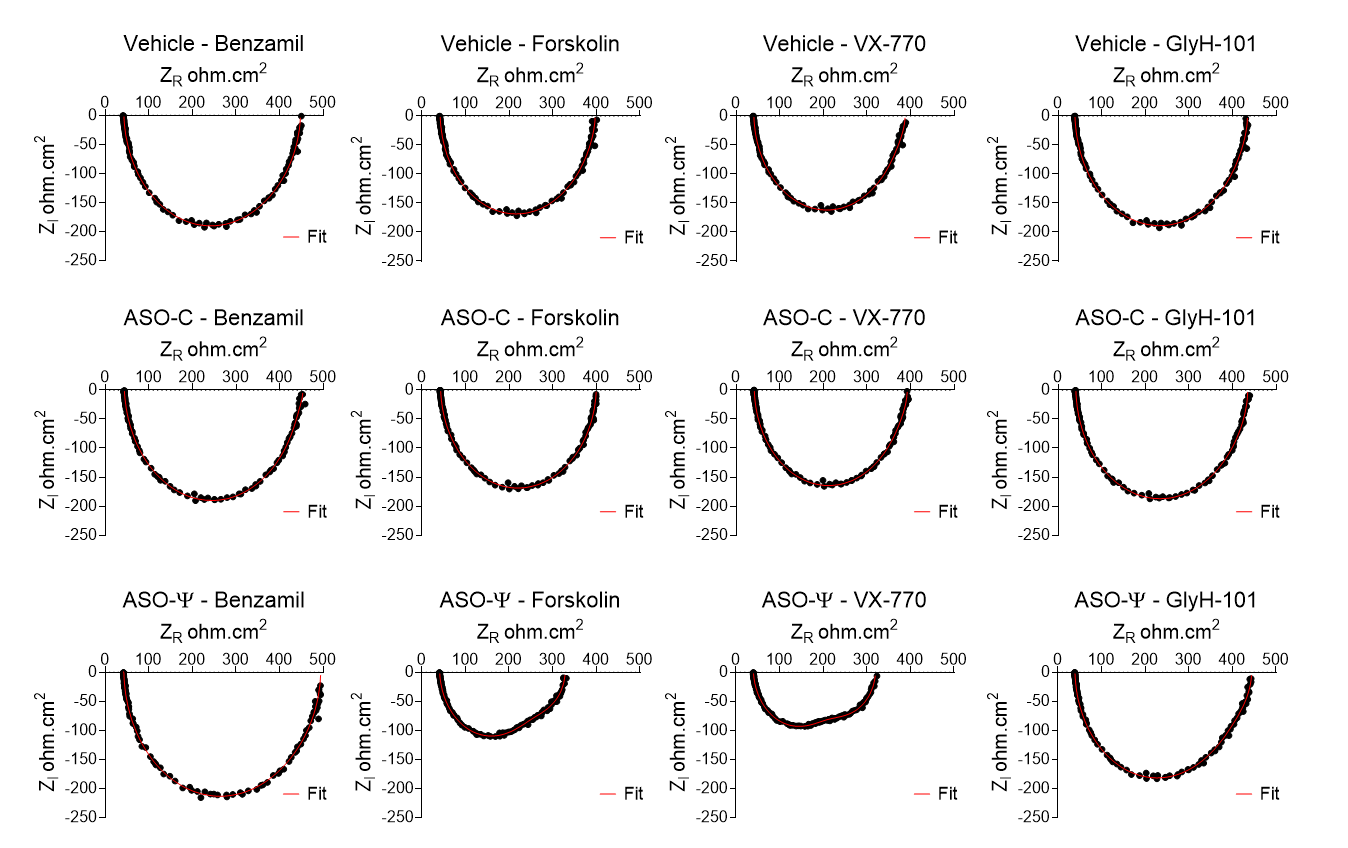

**Supplementary Figure S5.** Impedance spectra data presented as representative Nyquist plots from a single replicate after benzamil, forskolin, VX-770, and GlyH-101 treatments, in vehicle, ASO-C, or ASO-ψ treated homozygous *CFTR* c.3718-2477C>T hBE cells. Both the real (Z_R_) and imaginary (Z_i_) components of the measured impedance are plotted together from all 126 frequencies (black dots). Model fitting for each impedance spectrum is plotted as a red line (Fit).
