## Supplementary figures and images for "Antisense oligonucleotide-mediated correction of *CFTR* splicing improves chloride secretion in cystic fibrosis patient-derived bronchial epithelial cells"

### Supplementary Table S1

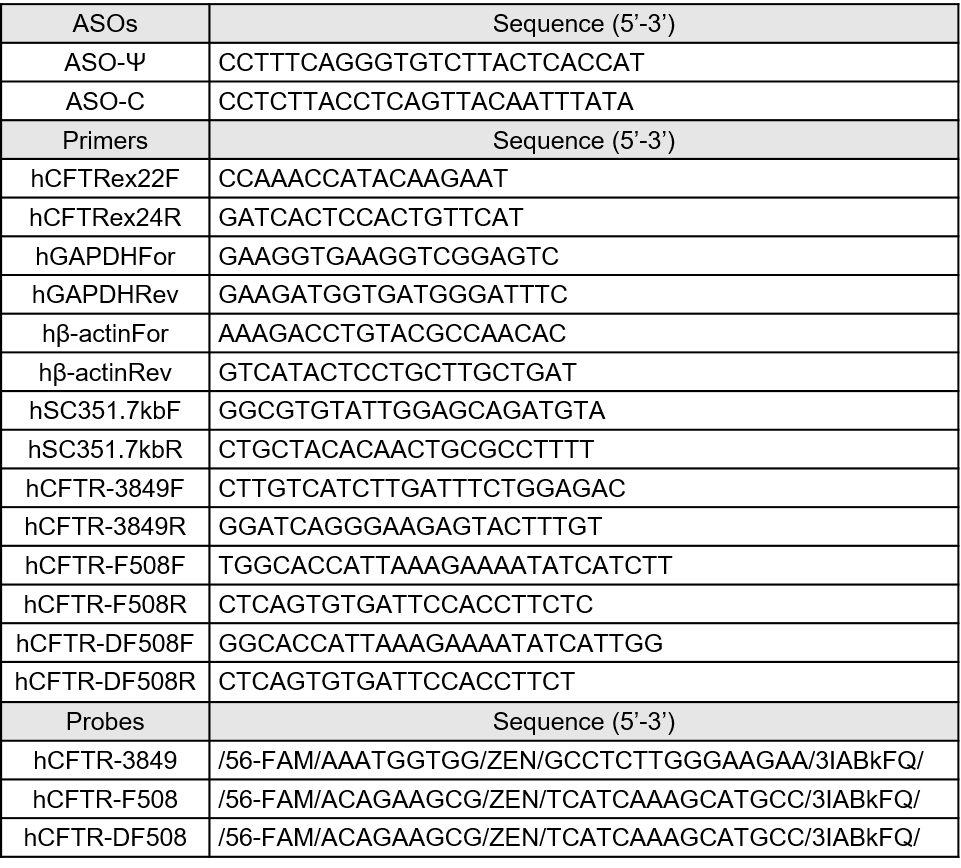

**Supplementary Table S1 –** Splice-switching Antisense Oligonucleotides, Primers, and Probes.

### Supplementary Table S3

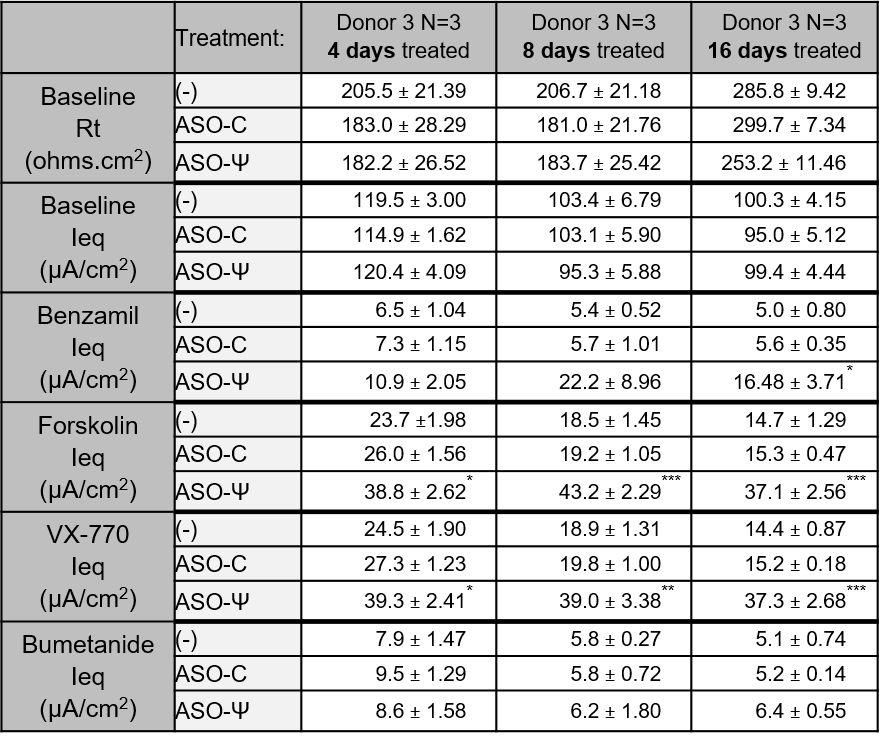

*=ASO-C vs ASO-Ψ	*p<.05, **p<.01, ***p<.001, ****p<.0001

**Supplementary Table S3.** I_eq_ and Rt values from donor 3 in Figure 4.
