## Supplementary Table S2 for "Antisense oligonucleotide-mediated correction of *CFTR* splicing improves chloride secretion in cystic fibrosis patient-derived bronchial epithelial cells"

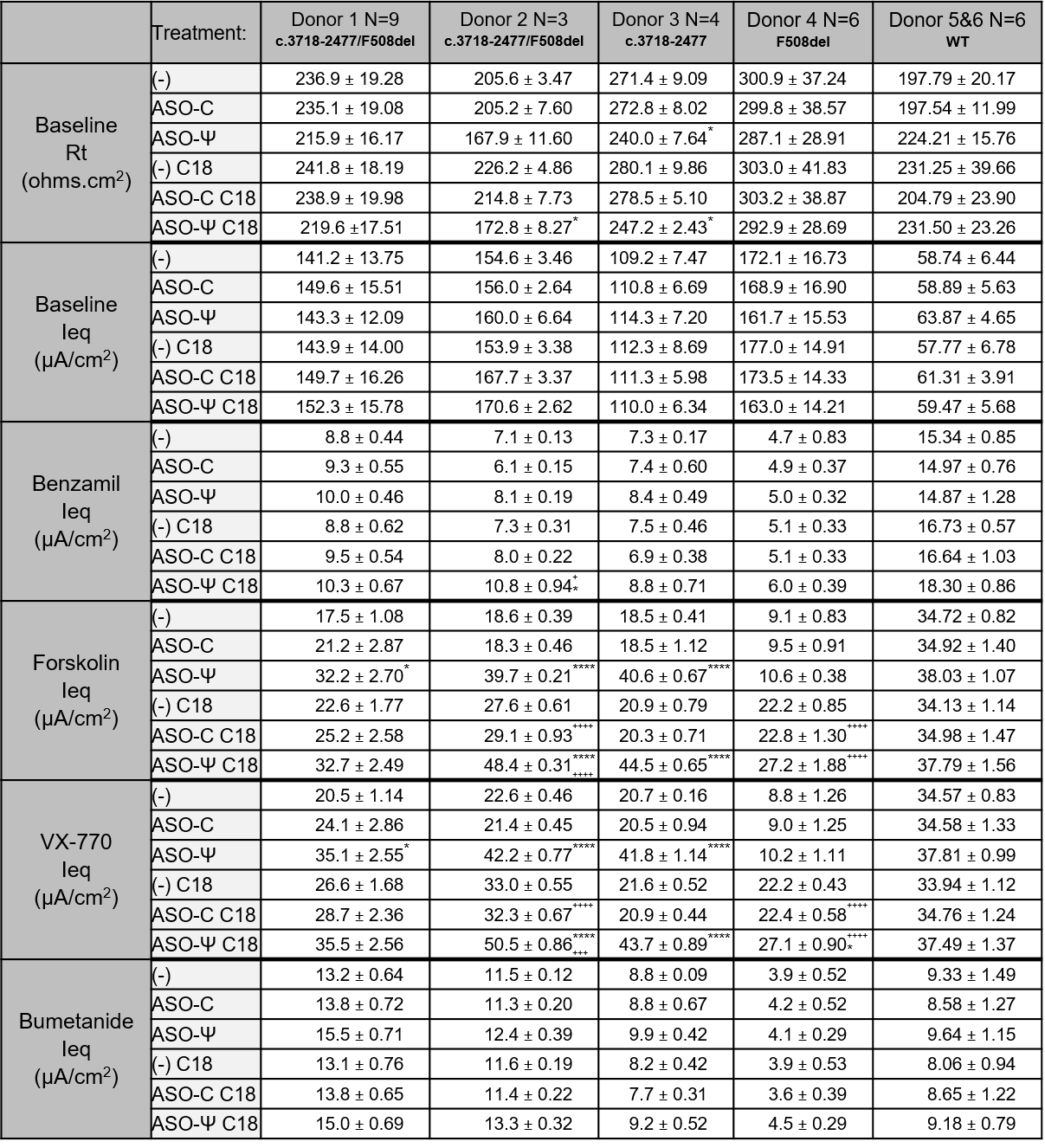

*=ASO-C vs ASO-Ψ	*p<.05, **p<.01, ***p<.001, ****p<.0001

+=ASO alone vs ASO + C18	^+^p<.05, ^++^p<.01, ^+++^p<.001, ^++++^p<.0001

**Supplementary Table S2.** I_eq_ and Rt values from donor HBE cell in Figures 2 and 6 and Supplementary Figures S2 and S3.
